# Diversity and Evolution of Computationally Predicted T Cell Epitopes against Human Respiratory Syncytial Virus

**DOI:** 10.1101/2022.06.29.498066

**Authors:** Jiani Chen, Swan Tan, Vasanthi Avadhanula, Leonard Moise, Pedro A Piedra, Anne S De Groot, Justin Bahl

## Abstract

Human respiratory syncytial virus (RSV) is a major cause of lower respiratory infection. Despite more than 60 years of research, there is no licensed vaccine. While B cell response is a major focus for vaccine design, the T cell epitope profile of RSV is also important for vaccine development. Here, we computationally predicted putative T cell epitopes in the Fusion protein (F) and Glycoprotein (G) of RSV wild circulating strains by predicting Major Histocompatibility Complex (MHC) class I and class II binding affinity. We limited our inferences to conserved epitopes in both F and G proteins that have been experimentally validated. We applied multidimensional scaling (MDS) to construct T cell epitope landscapes to investigate the diversity and evolution of T cell profiles across different RSV strains. We find the RSV strains are clustered into three RSV-A groups and two RSV-B groups on this T epitope landscape. These clusters represent divergent RSV strains with potentially different immunogenic profiles. In addition, our results show a greater proportion of F protein T cell epitope content conservation among recent epidemic strains, whereas the G protein T cell epitope content was decreased. Importantly, our results suggest that RSV-A and RSV-B have different patterns of epitope drift and replacement and that RSV-B vaccines may need more frequent updates. Our study provides a novel framework to study RSV T cell epitope evolution. Understanding the patterns of T cell epitope conservation and change may be valuable for vaccine design and assessment.

**Author Summary:** Lower respiratory infections caused by human respiratory syncytial virus (RSV) is a global health challenge. B cell epitope immune response has been the major focus of RSV vaccine and therapeutic development. However, T cell epitope induced immunity plays an important role in the resolution of RSV infection. While RSV genetic diversity has been widely reported, few studies focus on RSV T epitope diversity, which can influence vaccine effectiveness. Here, we use computationally predicted T cell epitope profiles of circulating strains to characterize the diversity and evolution of the T cell epitope of RSV A and B. We systematically evaluate the T epitope profile of RSV F and G proteins. We provide a T cell epitope landscape visualization that shows co-circulation of three RSV-A groups and two RSV-B groups, suggesting potentially distinct T cell immunity. Furthermore, our study shows different levels of F and G protein T cell epitope content conservation, which may be important to correlate with duration of vaccine protection. This study provides a novel framework to study RSV T cell epitope evolution, infer RSV T cell immunity at population levels and monitor RSV vaccine effectiveness.

## Introduction

Human respiratory syncytial virus (RSV) is a negative-strand RNA virus that is classified in the *Orthopneumovirus* genus of the family *Pneumoviridae*. It is a major cause of lower respiratory disease in young infants, immunocompromised individuals, and elderly people, resulting in annual epidemics worldwide [1]. The single-stranded RNA genome of RSV is approximate 15.2 kb and encodes 11 viral proteins [2]. The Fusion (F) and Glycoprotein (G) proteins are the two major surface proteins [3]. F protein is generally thought to be conserved and therefore it is the focus of most current RSV vaccine designs. Although G protein is highly variable, its contribution to disease pathogenesis and its role in the biology of infection suggest it can also be an effective RSV vaccine antigen [4]. Despite the significant burden of RSV infection worldwide, there is no licensed vaccine. The only approved intervention is passive immuno-prophylaxis with palivizumab, which is achieved by administering the monoclonal antibody (mAb) to a highly restricted group of infants under the age of 24 months and treatment must be repeated monthly during the RSV season due to the relatively short half-life of the antibody[5], [6]. Due to the high cost of monoclonal antibody treatments, this intervention is limited to high-risk infants and is generally unavailable in developing countries. An RSV vaccine is an urgent global healthcare priority, and it is likely that different strategies are needed for the various high-risk groups.

A number of research teams have worked on the development of RSV vaccine since its isolation and characterization in 1956 [7], [8]. However, vaccination with the formalin-inactivated, alum precipitated RSV (FI-RSV) vaccine in RSV-naïve infants and young children, led to the development of vaccine enhanced disease (VED) that hampered vaccine development for decades to follow [9]. Many studies have been conducted to explain this undesirable outcome. It is likely that formalin fixation led to a vaccine that mostly presented the post-fusion conformation of RSV F protein, leading to an excess of non-neutralizing antibodies and immune complex formation [10] [11] [12]. Other studies indicated that an impaired T cell response with Th2 skewing [13], [14], as well as complement deposition in the lungs, contributed to enhanced neutrophil recruitment [12]. Due to the recent breakthrough to structural constrain the F protein in the pre-fusion conformation and the development of RSV rodent models, there has been a surge in the number of RSV vaccine candidates undergoing clinical evaluation.

While most current RSV vaccination strategies focus on a B-cell-induced neutralization immune response, T cell immunity also plays a major role in the resolution of virus infection and is essential for RSV vaccine development [15], [16]. Once RSV infection of the lower airways is established, CD8 T cells play an important part in viral clearance and CD4 helper T cells can orchestrate cellular immune responses and stimulate B cells to produce antibodies. However, Th2-biased responses have been associated with animal models of RSV VED, and measurement of Th1 and Th2 responses are considered important to predict the safety of vaccine candidates [12]. Therefore, induction of a balanced cell-mediated immune response through vaccination would promote RSV clearance, but caution must be taken to avoid the potential for immunopathology. Taken together, a closer examination of T cell immunity and the virus sequences that induce T cell responses are needed for RSV vaccine development.

Human respiratory syncytial virus has a complex circulation pattern in the human population. Within two antigenic groups, RSV-A and RSV-B, different genotypes can co-circulate within the same community, while novel RSV genotypes with high genomic diversity may arise and potentially replace the previously dominant genotypes [17]. In recent years, several unique genetic modifications in RSV have been identified, including a 72-nucleotide (nt) duplication (ON genotype) in RSV-A G gene and another with a 60-nt duplication (BA genotype) in RSV-B at a similar region [18]. The observed RSV genetic diversity has raised a question about whether it is necessary for an RSV vaccine to include several different strains to be effective. Most current RSV vaccine developments are based on an RSV A2 laboratory strain, which is a chimeric strain that belongs to subtype A [19]. While these treatments hold promise, there is the possibility of viral strains developing escape mutations. For example, palivizumab-resistant strains have been isolated from both RSV rodent models and human [20][16]. Monoclonal antibody tests have demonstrated additional antigenic variability within RSV-A /B antigenic groups and suggest that it may play a role in the ability of RSV to escape immune response and established infections [21]. In addition, amino-acid variation at the T cell epitope level and the emergence of novel T cell epitopes have been reported [22], but further studies are needed to illustrate the effect of these variations on T cell recognition. T cell epitopes are sometimes cross-reactive, which is defined as the recognition of two or more epitope peptide-MHC complexes by the same T cell receptor and these cross-reactive epitopes are predicted to be cross-conserved [23]. Prediction of cross-conservation is important for vaccine design because it would be useful to predict protection against a pathogen with different lineages and identify escape variants. Hence, characterizing T cell epitope profiles across different strains can be crucial for RSV vaccine development.

In this study, we utilize immunoinformatic approaches that are implemented in the iVAX toolkit [24] to predict T cell epitopes in RSV across different strains with a focus on the two major surface proteins F and G. With the analysis of a comprehensive dataset, we evaluate the lineage-specific T cell epitope profile of RSV. We also create sequence-based T cell epitope landscapes based on epitope content comparison across different strains and further correlate RSV T cell immunity change with virus evolution. The proportion of cross conserved T cell epitope content between vaccine candidate strains that developed earlier and RSV circulating strains with different isolated years and locations were also calculated. These analyses may aid in understanding RSV T cell immunity across different strains and contribute to current vaccine design efforts.

## Results

### Distribution of T cell epitopes in RSV surface proteins

We evaluated the T cell immunogenic potential across RSV surface proteins by scanning 9 residue regions to predict the binding probability to MHC class I and class II molecules (Figure 1). The epitope density of RSV surface proteins was evaluated using a normalized epitope density score. F protein has an epitope density score greater than +20 for both the class I and class II immunogenicity scale analysis, indicating significant immunogenic potential [24]. This contrasts with lower G protein class I and class II epitope density protein scores for both subtypes. The class I epitope density score of G protein was greater than +10 in both subtypes but the class II density was lower than random expectation in the analysis of RSV-B (Figure 1A). This result suggests that RSV surface proteins are likely to have the potential to stimulate T cells that are required for protective immunity. We then investigated the distribution of T cell immunogenicity across the proteins and found that there are regions with relatively high T cell immunogenic potential (Figure 1B). The distribution of T cell immunogenicity of F protein was mapped onto its protein structure and overlap between protein sequence regions with high T cell immunity potential and the antibody neutralizing targets was observed at antigenic site Φ and site II.

**Figure 1:**
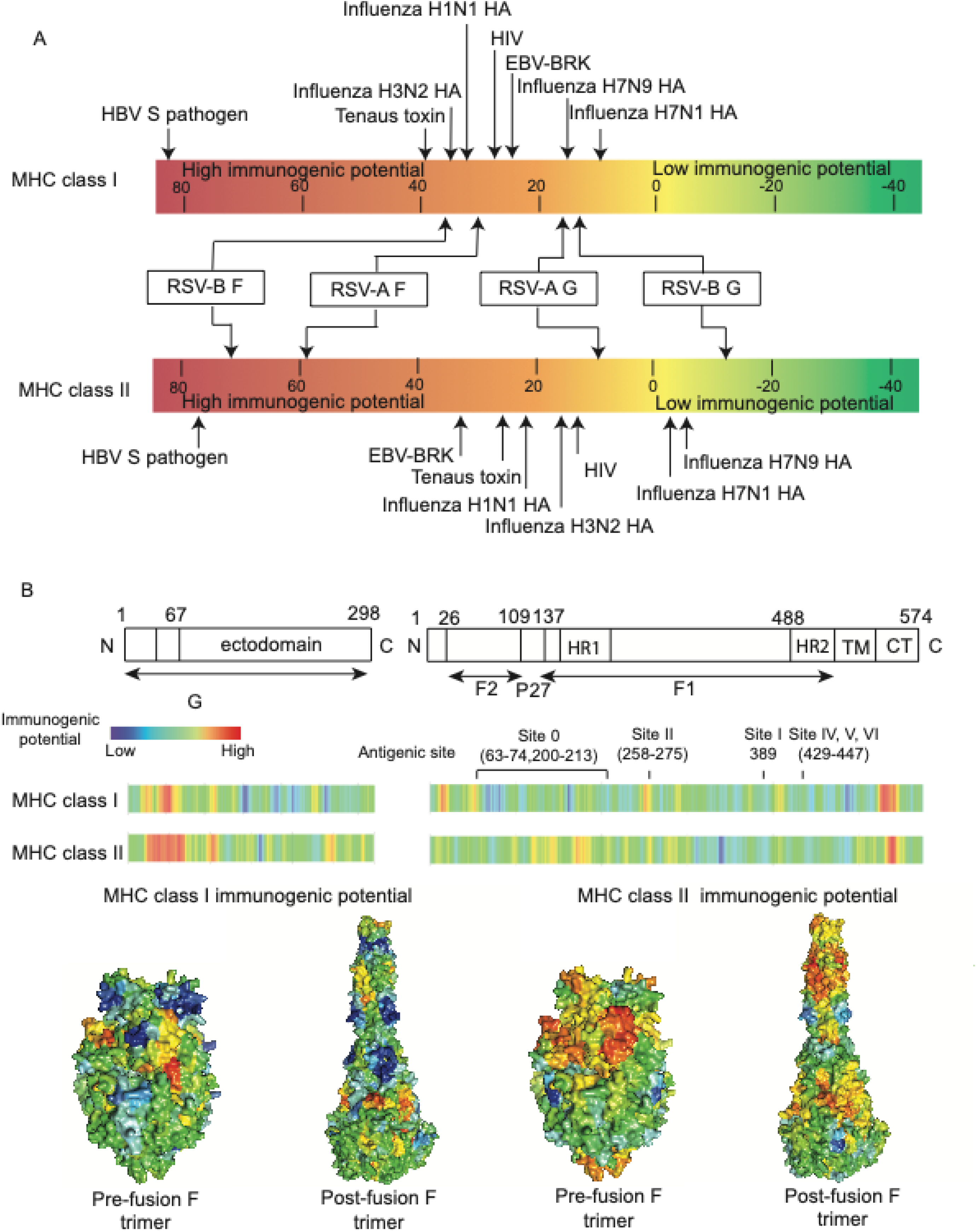
T cell immunogenic potential for RSV surface proteins based on MHC binding prediction. (A) T cell immunogenic potential of RSV major surface proteins. T cell epitope density scores for RSV major surface proteins and other pathogen proteins are labeled on a scale bar. Low-scoring proteins are known to engender little to no immunogenicity while higher-scoring proteins are known immunogens. Proteins scoring above +20 on this scale are considered to have significant immunogenic potential. (B) Distribution of RSV T cell immunogenic potential across F and G protein in RSV reference strain A2. Prefusion or post-fusion F protein surface was colored by the relative immunogenetic potential at each location. Analyses are based on the RSV-A reference sequence.

### Lineage specific T cell epitope profiles

We then extended T cell epitope predictions from RSV representative strains to multiple wild-circulating strains. The distribution and diversity of T cell epitopes across different strains are illustrated in heatmaps with the corresponding time-scaled phylogenies (Supplementary Figure 1 and Supplementary Figure 2). Both F and G proteins contain epitopes that were conserved across all RSV strains in almost 100% of sampled isolates, suggesting that they could serve as high-quality T cell epitope candidates for vaccine design. In contrast, some epitopes were mutated in selected strains, and those epitopes that only occurred in certain clades within the phylogeny could be interpreted as clade-specific “fingerprints”.

The G gene duplication events in RSV, which are unique gene signatures, can either shift the position of epitopes or cause the emergence of novel epitopes. Two novel class I epitopes, (no. 31 and no. 40 in Supplementary Figure 2A), were found in RSV-A strains that contain G gene duplication. In addition, an emergent class II epitope (no. 25 in Supplementary Figure 2A) was identified in RSV-A sequences that contain G gene duplication, which was a shift from an epitope (no. 24) that has been observed in other strains. From RSV-B strains that contain the G gene duplication event, we also observed multiple lineage specific class I T cell epitopes, which are caused by a 2-aa deletion (aa157 and aa158) in these strains instead of directly due to the 60-nt duplication event. RSV-B G proteins that have the duplication event contain multiple novel epitopes (no. 22, 23, 26, 28, 30, 37) but do not contain several epitopes (no. 24, 25, 27, 29, 31, 38) that are identified in other strains (Supplementary Figure 2B).

To further determine whether the T cell epitopes defined using EpiMatrix might be immunogenic, we utilized the JanusMatrix [25] algorithm to identify the T cell epitopes that are likely to be cross-conserved with human epitopes and thereby tolerated by the immune system. Based on this analysis, 6.45% of putative class I epitopes and 1.12 % of putative class II epitopes of RSV major surface proteins are cross-conserved with human proteome-derived epitopes at TCR facing residues. As these peptides have similar HLA binding preferences that are contained in human proteins (Supplementary Figure 3), they were therefore assumed not to be immunogenic. After excluding the high-JanusMatrix score epitopes identified above, we were able to identify T cell epitopes that were conserved in more than 60% of currently circulating RSV strains. We searched the IEDB epitope database to determine if these epitopes were related to experimentally validated RSV T cell epitopes or HLA ligands. The conserved RSV T cell epitope sequences that may be important for future vaccine development are shown in Table 1 and Table 2.

**Table 1:**
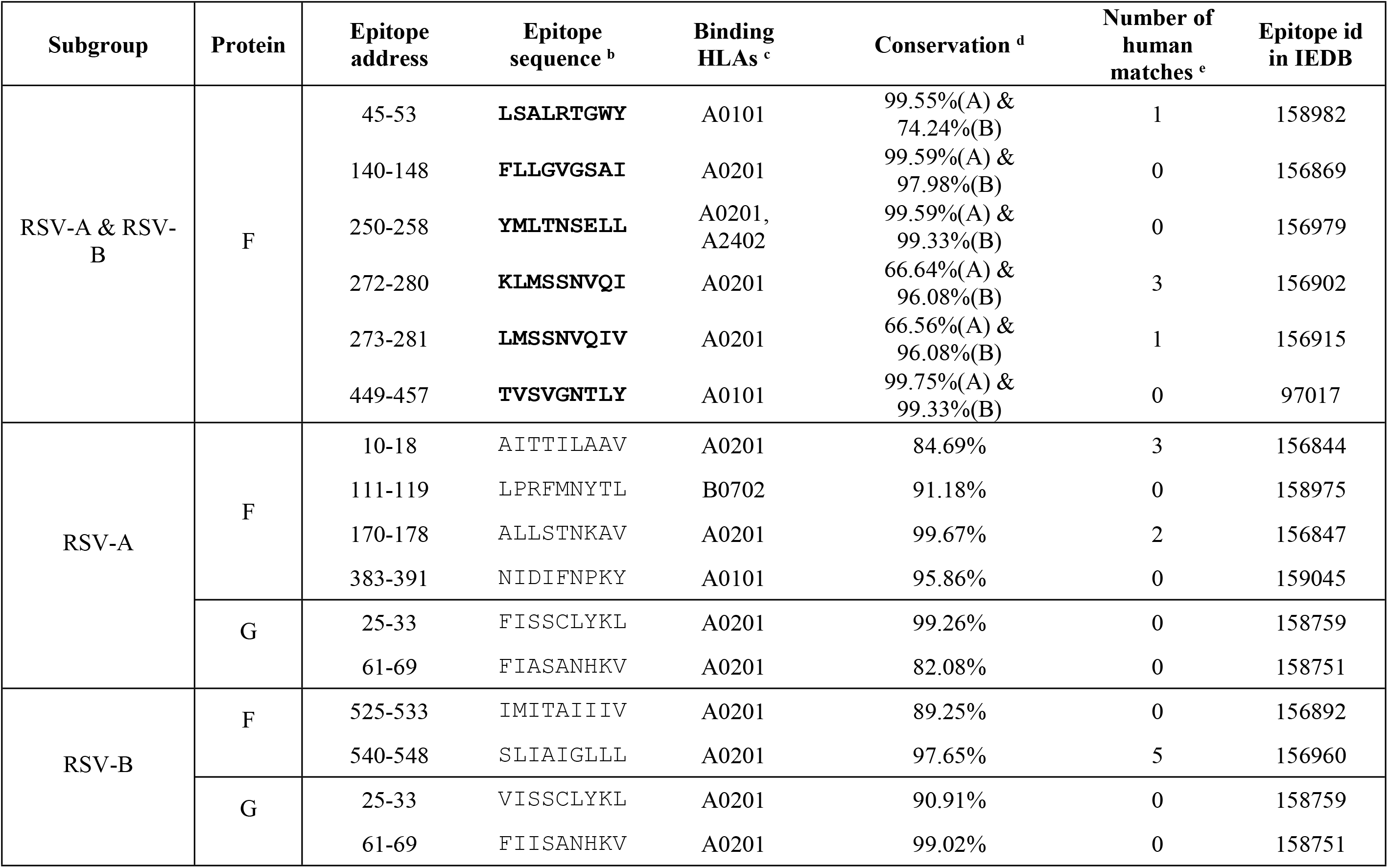
Experimentally validated conserved MHC class I epitopes peptides in RSV major surface proteins. a. This table contains putative MHC class I epitopes that have already been experimentally validated in publications. b. Epitopes sequences that are conserved in both RSV-A and RSV-B are in bold. c. HLAs that have the top 1% binder scores in EpiMatrix for epitope sequence. d. The conservation is evaluated by the presence of epitope peptides across all RSV-A or RSV-B sequences that are publicly available (only epitope sequences with at least 60% conservation are shown in the table). e. Count of human epitopes found in the search database. JanusMatrix was used to search human epitopes that are predicted to bind to the same allele as the RSV epitope and share TCR facing contacts with the RSV epitope.

**Table 2:**
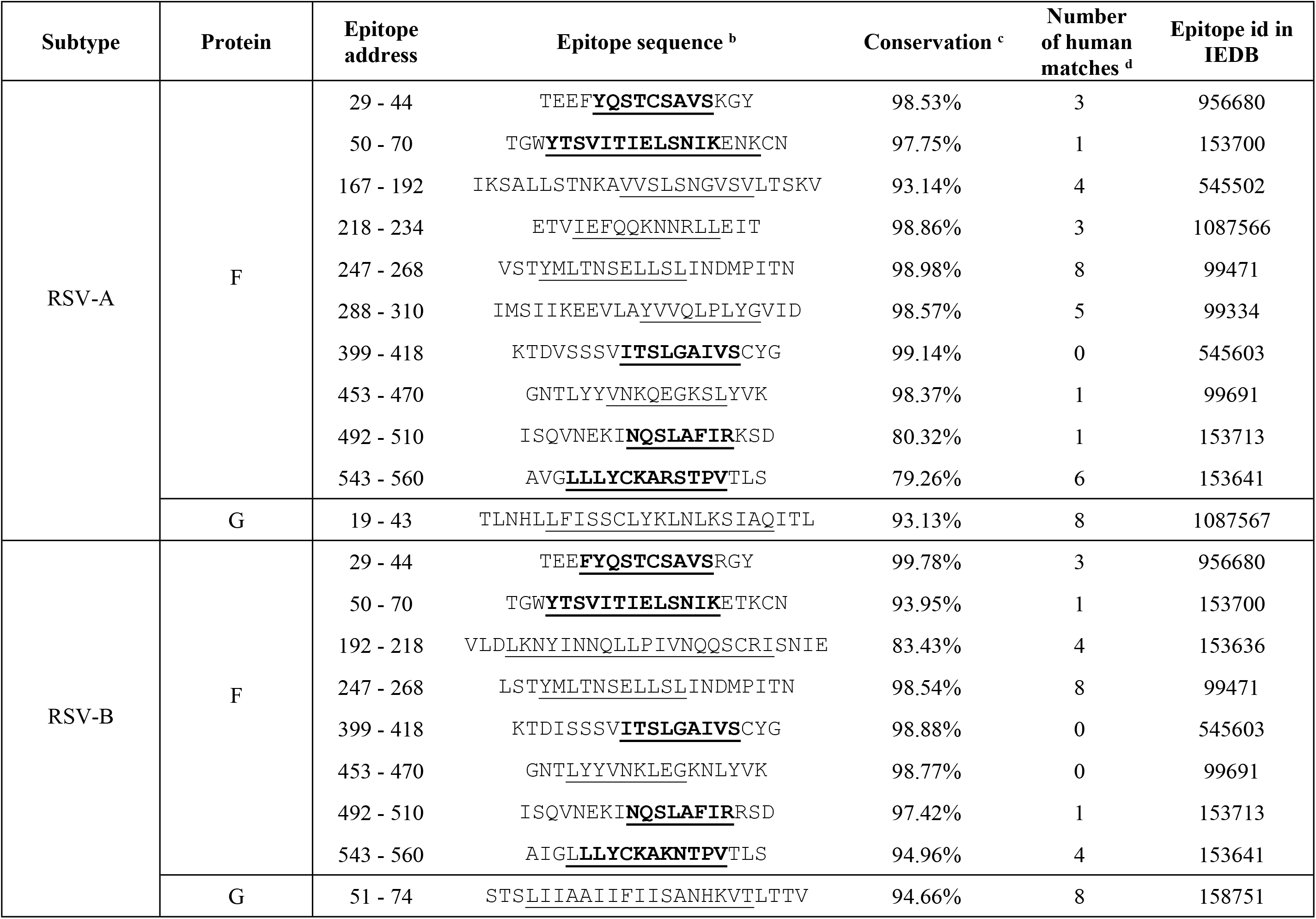
Experimentally validated conserved MHC class II epitopes peptides in RSV major surface proteins. a. This table contains putative MHC class II epitopes which share the identical binding groove sequence of the RSV class II epitopes that have already been experimentally validated in publications. b. Underlined sequences represent the nine-mer frames with the greatest potential to bind class II HLA. Epitope sequences that are in bold indicate sequences are predicted to bind class II HLA and are conserved in both RSV-A and RSV-B. c. Conservation is evaluated by the presence of epitope peptides across all RSV-A or RSV-B sequences that are publicly available (Only epitope sequences with at least 60% conservation are shown in the table). d. Count of human epitopes found in the search database. JanusMatrix was used to search human epitopes that are predicted to bind to the same allele as the RSV epitope and share TCR facing contacts with the RSV epitope.

### Predicted RSV T cell epitope landscapes

To investigate the evolution of RSV on T cell immunity profiles, we develop a new approach to visualize the T immunity profile of multiple RSV strains on a landscape. We performed a T cell epitope content pairwise comparison between RSV strains, and T cell epitope distances between pair of RSV strains were then calculated using a T epitope distance algorithm. We then applied a multidimensional scaling (MDS) approach using these estimated pair-wise T epitope distances to map RSV strains to a landscape to characterize their T-cell immunity profile. We found both Class I and Class II T cell immunity profiles of F and G proteins of different RSV strains were clustered into groups on this T cell epitope landscapes (Supplementary Figure 4). Combining the Class I and Class II T-cell epitope biding profiles, RSV-A major surface protein isolates can be divided into three clusters and RSV-B major surface protein isolates can be divided into two clusters (Figure 2, Supplementary Figure 5). We observe that the G gene sequence isolates that contain 72-nt (RSV-A) or 60-nt (RSV-B) duplications clustered together with other sequences instead of forming isolated groups. To further investigate the T cell epitope diversity, we correlated this clustering pattern with the phylogenetic histories (Figure 2B). The phylogenetic tree topologies of the RSV-A F gene and G gene are similar. The F gene cluster 1 is paraphyletic, while cluster 2 and 3 are monophyletic. Cluster 1 is the closest to the ancestral sequence and mapping this group onto the phylogeny show that this cluster has a basal relationship with clusters 2 and 3 indicating that the phylogenetic divergence occurred prior to epitope drift. The RSV-B F and G gene genealogies are very different. In particular, the RSV-B F gene topologies is indicative of strong immune selection, similar to observed human influenza A virus or within host HIV phylogenies [26]. In contrast, the RSV-B G gene phylogeny shows the co-circulation of multiple lineages, though this could reflect the sequencing bias of G genes (Figure 2B). We then calculated the T-cell epitope immune distance of each strain from a reconstructed ancestral sequence (Figure 2C). These distances were then plotted against the year of isolation and colored according to the cluster identified in Figure 2A. RSV-A shows that multiple predicted immune phenotypes co-circulate and persist for long periods (>2 decades). Analysis of RSV-B shows a turnover of the predicted immune phenotypes with short periods of co-circulation (<5 years) for F and G protein T cell epitopes. The limited periods of co-circulation is again consistent with phenotype patterns observed for viruses under strong immune selection (e.g H3N2 influenza A virus) [27], [28]. In contrast, genetic distances from the reconstructed ancestral sequence plotted against year of isolation show patterns typical of gradual genetic drift, except in the G gene where a 72-nt and 60-nt insertion is present (Figure 2D). Taken together, these results suggest that genetic and predicted T-cell epitope immune diversity are different and may be an important factor to consider when evaluating RSV vaccine efficacy.

**Figure 2:**
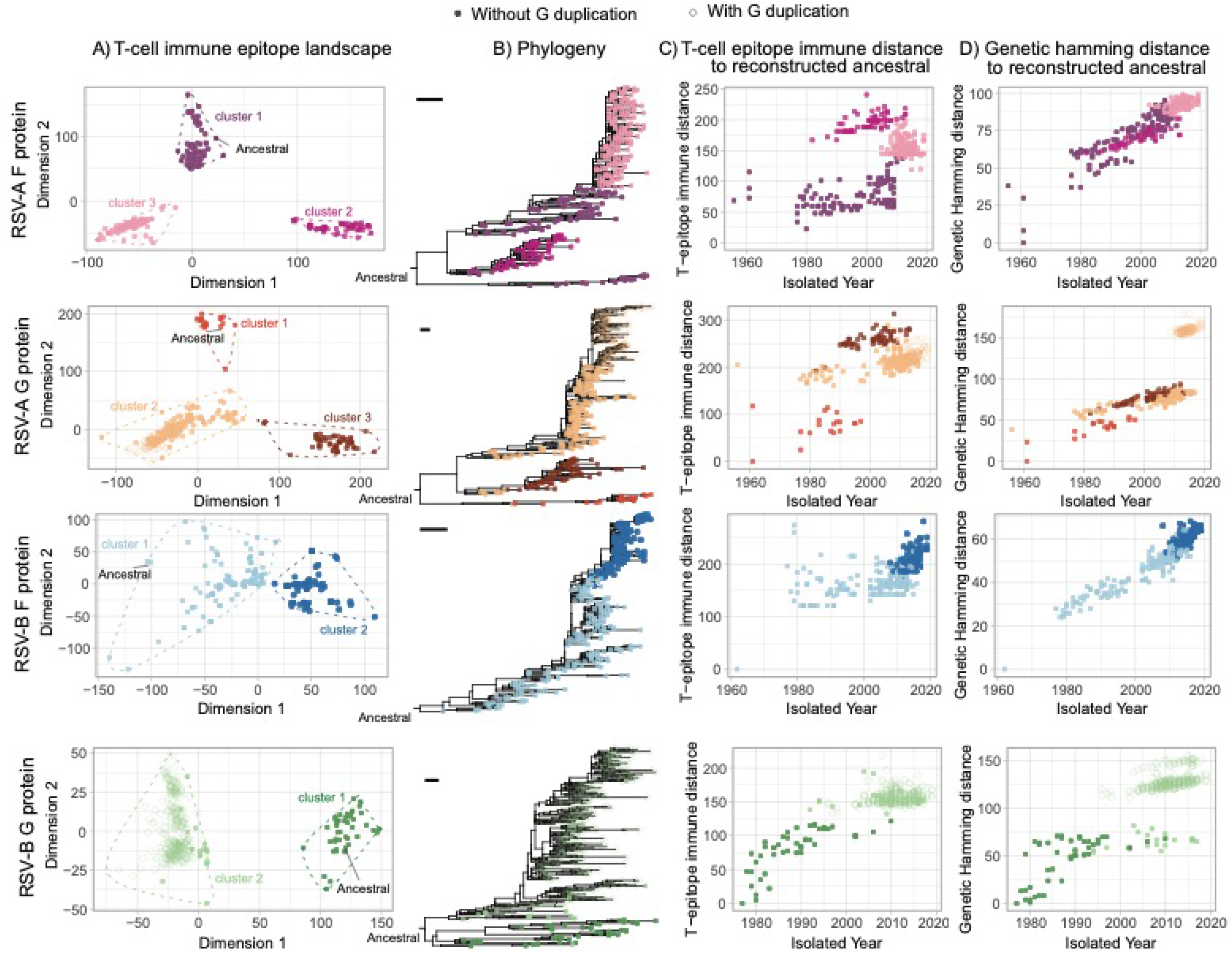
Predicted T cell epitope landscapes and genetic evolution of RSV surface proteins. Epitope landscapes of RSV major surface proteins are built with MHC class I and class II epitope content comparison across different strains. Filled circles indicate F protein isolates or G protein isolates without duplication. Diamonds indicate G protein isolates with gene duplication. T cell immunity clusters are determined with *k-means* method and are used to color the sequenced isolates in the following panels. The corresponding Maximum Likelihood (ML) phylogenies are reconstructed and are rooted by mid-point. Scale bars indicate 0.005 nucleotide substitution per site. T cell epitope immune distance and genetic hamming distance from the estimated TMRCA are plotted against the isolated time of each sequence.

There are multiple methods available to predict T cell epitopes [29], which may result in different reconstructed landscapes if there is a systematic bias in the prediction method. We used the NetMHCpan method [30] to predict T cell epitopes and perform the same landscape reconstruction using MHC class I binding predictions for RSV-A F protein. Our analysis showed a consistent clustered pattern of RSV T epitope profile on the landscape regardless of T cell epitope prediction method (Supplementary Figure 6).

### Assessment of vaccine candidate strains with T cell epitope content

To quantitatively evaluate whether it might be necessary to include multiple RSV strains to prepare an effective vaccine, two live attenuated RSV strains that are previously considered as vaccine candidates, CP248, a recombinant virus that belongs to subtype A, and CP52, which is a recombinant RSV-B strain, were included in our analysis and compared to wild-type strains using EpiCC. We calculated the average proportion of cross-conserved T cell epitope content between the selected vaccine strains and wild-circulating strains from different isolation years and WHO regional groups (Figure 3, Supplementary Figure 7). Different proportions of cross-conserved T cell epitope content against isolates from two different subtypes, A and B, were observed in both the F and G protein analyses. In the comparison of the vaccine strains and wild strains belonging to the same subtype, the proportion of cross-conserved T cell epitope in RSV F protein is relatively stable in different groups, all are higher than 78% for RSV-A and higher than 85% for RSV-B. In contrast, changes in the proportion of cross-conserved T cell epitopes were detected among groups within the same RSV subtype, especially in different temporal groups in the G protein analysis (Figure 3). Vaccine strain CP248 appears to have a relatively higher proportion of cross conserved T cell epitopes within G protein when compared to the RSV-A strains that were isolated before 1991 (> 70%) and a relatively lower degree of conservation against recently isolated strains. A similar decrease in T cell epitope conservation with time was identified for vaccine strain CP52 among circulating RSV-B strains.

**Figure 3:**
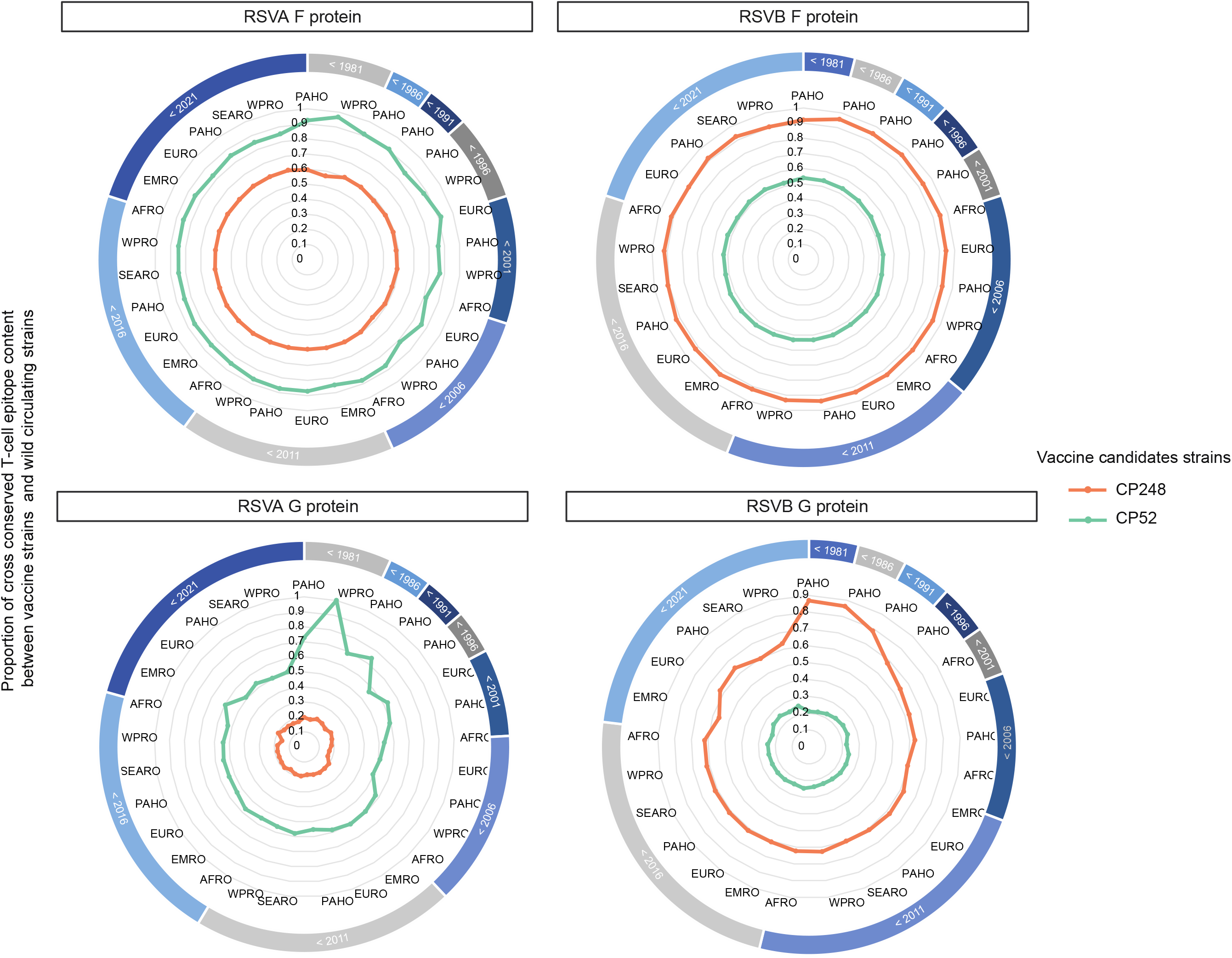
Evaluation of previously used RSV vaccine candidate strains with T cell epitope content of circulating strains. RSV-A and RSV-B major surface protein sequences are subsampled and then grouped by isolation year and 6 isolated WHO regions. African Region (AFRO), Region of the Americas (PAHO), South-East Asia Region (SEARO), European Region (EURO), Eastern Mediterranean Region (EMRO) and Western Pacific Region (WPRO). The proportion of cross-conserved T cell epitope content between live attenuated strains (CP248 or CP52) and wild circulating strains are displayed as radar plots.

## Discussion

Although both CD4 and CD8 T cells contribute to protection against RSV-induced disease following primary infection [16][31], T cell epitopes have received limited attention in the RSV research effort. We demonstrate RSV surface proteins appear to have significant potential to drive T cell immunity using a computational approach, based on their T cell epitope density scores as determined by MHC molecular binding prediction. The relatively high putative T-cell epitope density might make F protein a good target for RSV vaccine. In addition to the analysis of T cell epitope density and distribution in RSV major surface proteins, we also demonstrated lineage-specific variations in T cell epitope content. Even though RSV F protein is believed to be well conserved and G protein is reported to be highly variable, epitope mutations are observed across different lineages within the F protein, and potential conserved T cell epitopes can still be found in the highly variable G protein, suggesting that studying the lineage-specific T cell epitopes in RSV can provide insight into the impact of immune selection on viral diversity and persistence. While experimental validation is needed, this analysis highlights the importance of understanding population-level epitope conservation as it may provide important insight into the development of T cell epitope-driven vaccines against RSV infection.

A major focus of our work is the development of a sequence-based method to map the evolution of T cell immunity across different stains. Following a previously pivotal work that used MDS method to map the evolutionary adaptation of influenza A virus-induced by CD8 T cell using the presence and absence of MHC class I epitopes [32], we constructed RSV T cell immunity landscapes using immune distances that were generated by T cell epitope cross-conservation analyses, which allows for easy visualization and intuitive understanding of the potential for T cell immunity relationships among different strains. When comparing across strains, we found that the T cell epitope content of RSV surface proteins from different strains can be clustered, as has been observed for the antigenic relationship reported in other pathogens [49][50]. Our results also demonstrate the correspondence between RSV T cell immunity clusters and their corresponding phylogeny, with sequences in the same clade generally belonging to the same T cell immune cluster. Importantly, we also observe different patterns of T-cell epitope evolution of RSV wild strains compared with their genetic evolution, which highlights the importance of characterizing T cell epitope changes in RSV.

We identified highly conserved RSV T cell epitopes in this study, some of which have already been experimentally validated and published in the IEDB database. However, we also identified several other conserved T cell epitopes that have not been previously described. These may be valuable for vaccine design, although experimental validation will be needed. Furthermore, the homology of selected RSV epitopes to human epitopes suggests that some predicted RSV T cell epitopes might be tolerated by the human immune system, or could induce a harmful cross-reactive immune response against human proteins when administered with an adjuvant [25]. Certain aspects of immunity to RSV were not addressed by this study. For example, neutralizing antibody responses are currently considered to be the most important correlate of immunity. While neutralizing antibodies would not directly be elicited by a T cell epitope-driven vaccine, helper (CD4) T cell epitopes are required to generate high affinity, high specificity antibodies. We also note that we have limited our focus on the two major RSV surface proteins in our current analysis, but other RSV proteins like N, M, or M2-2 proteins might also contribute to vaccine efficacy [36].

An effective vaccine against variable viruses should contain T cell epitopes that are highly conserved among circulating strains [37]. Vaccine efficacy can be diminished if T cell epitopes in a vaccine strain do not match when new strains of pathogens emerge. In this study, we used an immunoinformatic-based approach to estimate cross-conserved T cell epitope contents between two live attenuated vaccine candidate strains and RSV circulating wild strains. We found that there was a low proportion of cross-conserved T cell epitope content with vaccine strains that belonged to different antigenic groups, which indicates the risk of using a single-subtype strain in RSV vaccines. In addition, we observed a lower proportion of cross-conserved G-protein T cell epitope content between vaccine strains and recent circulating strains in the same antigenic group, which suggests that including T cell epitopes from different strains in the same antigenic group might also be important for RSV vaccine development. Although we did not observe a significant change in cross-conserved T cell epitope content in F protein, we cannot rule out the probability that variation of F protein in the future could render a single-strain-based vaccine less effective. Our current analysis is based on reduced datasets due to the heavy computational capacity required to perform epitope content comparison. We constructed these representative datasets by randomly subsampling the complete datasets according to geographical regions and isolated years. Our findings may reflect the T cell epitope diversity of publicly available RSV strains, however, additional RSV surveillance efforts may be required to get a full picture of the T cell epitope variability of RSV.

The lack of experimental data might cause problems in epitope identification. Computational-based T cell epitope landscapes have the potential for bias. But the observed clustered T cell evolutionary pattern in RSV surface proteins provide valuable insights into virus evolution in the aspects of T cell immunity and strain selection for vaccine design.

Overall, this study provides a focused analysis of T cell epitopes in RSV major surface proteins using computational tools. We performed a comprehensive T cell epitope prediction for RSV showing the immunological relationship of T cell epitopes in RSV surface proteins. This study demonstrates that T cell epitope evolution may differ from genetic variation and provides a framework for developing an integrated epitope-based RSV vaccine and evaluation methods that could be used to optimize vaccination strategies.

## Materials and Methods

### Dataset

RSV GenBank records files were retrieved from NCBI’s GenBank nucleotide database using the search term “HRSVA” or “HRSVB” on June 22, 2020. F and G gene nucleotide sequences and metadata including country of isolation and collection date were extracted using customized python scripts. Genotype assignments were made with the program “LABEL”, using a customized RSV module [38] [39]. Countries of isolation were grouped into 6 WHO regions: African Region, Region of the Americas, South-East Asia Region, European Region, Eastern Mediterranean Region, and Western Pacific Region [40]. The following inclusion and exclusion criteria were applied: (i) each sequence needed to have a known isolated geographic location and isolated year, (ii) each sequence had to be at least 80% of the complete gene sequence in length, (iii) identical sequences with the same isolate country were removed, and (iv) vaccine derivative and recombinant sequences were removed. Using these criteria, comprehensive datasets of RSV F and G genes were defined (RSV-A F gene = 1010, RSV-B F gene = 894, RSV-A G gene = 1488, RSV-B G gene = 1120). Nucleotide sequences from each dataset were aligned using MAFFT.v7 [41] and were translated into amino acids using EMBOSS.v6.6.0 [42] for immunoinformatic analyses. In addition, two artificial sequences, CP248 and CP52 (cold passage live RSV strains that were previously evaluated as vaccine candidates, Accession No: U63644, AF0132551 respectively) were downloaded from the NCBI’s GenBank nucleotide database [43].

### Phylogenetic inference

The nucleotide sequences of RSV major surface proteins were used to reconstruct the maximum-likelihood (ML) phylogeny of RSV using RAxML.v8 with GTR+GAMMA substitution model [44]. The best-scoring ML tree was automatically generated from five runs by RAxML. Time-scaled phylogenies were further reconstructed with the best scoring ML trees using the program “Timetree” [45]. The phylogenies are visualized in the R package “ggtree” [46].

### T cell epitope prediction

RSV major surface protein sequences were scored for binding potential against a globally representative panel of Human Leukocyte Antigen (HLA) class I and class II alleles using the EpiMatrix algorithm. This algorithm as well as the ClustiMer, JanusMatrix, and EpiCC algorithms discussed below are part of the iVAX toolkit developed by EpiVax, which is available for use under a license or through academic collaborations [24].

Evaluation of class I epitopes was made based on predictions for four HLA-A and two HLA-B supertype alleles: A*01:01, A*02:01, A*03:01, A*24:02, B*07:02, B*44:03. Class II epitopes were identified for nine HLA-DR supertype alleles: DRB1*01:01, DRB1*03:01, DRB1*04:01, DRB1*07:01, DRB1*08:01, DRB1*09:01, DRB1*11:01, DRB1*13:01, and DRB1*15:01. EpiMatrix parsed 9-mer sequence frames (each one overlapping the previous one by one amino acid) from the antigen sequence and assigned a score for each nine-mer/allele pair on a normalized Z distribution. Nine-mer sequences that had Z-scores of at least 1.64 are considered to be in the top 5% of any randomly generated set of 9-mer sequences and to have a high likelihood of binding to HLA molecules and being presented to T cells. Sequences that score above 2.32 on the Z-scale (top 1%) are extremely likely to bind to a particular HLA allele and to be immunogenic. For this analysis, HLA-class I restricted 9-mer sequences that had top 1% binder scores to at least one HLA class I supertype allele were considered to be putative class I epitopes [24].

To identify putative class II epitopes, we used an algorithm called ClustiMer [24] to screen EpiMatrix scoring results for the nine class II alleles. ClustiMer identifies contiguous regions of 15–30 amino acids that have a high density of MHC class II binding potential. Epitope density within a cluster is reported as an EpiMatrix Cluster Score, where scores of 10 and above are likely to be recognized in the context of multiple class II alleles and to be high-quality class II epitopes.

We also applied analysis of human homology to this study. The JanusMatrix algorithm was used to assess the potential cross-conservation of T cell epitopes with epitopes restricted by the same HLA alleles in the human proteome [25]. Briefly, JanusMatrix scans each identified epitope and examines the shared T cell receptor (TCR) contacts with class II epitopes present in the human proteome, to compute a JanusMatrix Human Homology Score. As defined in retrospective studies, foreign class I epitopes that score greater than 2 and class II epitopes that score greater than 5 may be less immunogenic due to T cell tolerance.

### Protein-level T cell immunogenic potential evaluation

RSV reference sequences (RSV-A: NC_038235, RSV-B: NC_001781) were downloaded from the NCBI RefSeq database and were used to evaluate the protein-level immunogenic potential of RSV major surface proteins. The protein-level immunogenic potential as represented by the EpiMatrix-defined T cell epitope density score was computed by summing the top 5% binder scores across HLA alleles and normalizing for a 1000-amino acid protein length. Zero on this scale is set to indicate the average number of top 5% binders that would be observed in 10,000 random protein sequences with natural amino acid frequencies. Proteins scoring above +20 have been observed to have the significant immunogenic potential [47]. Fully human proteins generally score lower than zero on the EpiMatrix immunogenicity scale.

To investigate the distribution of T cell immunogenic potential across RSV protein sequence regions, we summed up the binding scores of HLA alleles for each nine-mer frame, to get a frame-specific immunogenic potential score and standardized this score to a relative scale. The relative immunogenic potential across protein structure was represented by a color scale and the visualization of F protein structure was built with PyMOL Molecular Graphics System, Version 2.0 (Schrödinger, LLC). Protein data bank (PDB) files 5UDE [48] and 3RRR [49] were used for the pre-fusion and post-fusion forms.

### Subsampling strategy

Considering the heavy computational load that would be required to evaluate all available RSV sequences and to correct the overrepresentation of recently sampled strains, the comparative analysis for T cell epitope content was conducted with datasets in which overrepresented groups were reduced. A maximum of five sequences of each isolation year from different WHO region groups were subsampled randomly from the original datasets (RSV-A F gene = 402, RSV-B F gene = 319, RSV-A G gene = 390, RSV-B G gene = 359).

### T cell epitope content comparison

The Epitope Content Comparison (EpiCC) algorithm, which is implemented in iVAX was used to evaluate pairwise T cell epitope cross-conservation potential within each subsampled dataset [50]. Briefly, T cell epitope cross-conservation was evaluated by the binding likelihood of epitopes from different antigens with identical T cell receptor-facing residues (TCR*f*), which are predicted to bind to the same MHC allele. Two epitope sequences are assumed to be potentially cross-conserved if they have identical TCR*f* (position 4, 5, 6, 7, 8 for class I epitopes binding core and 2, 3, 5, 7, 8 for class II epitopes binding core) regardless of differences on their MHC-facing amino acids. To simplify the analysis, the binding of 9-mer epitopes within protein sequences are assumed to be mutually exclusive and uniform.

Therefore, T cell epitope immune distance (*D*) between two wild circulating strains (*w*_1_ and *w*_2_) can be defined as the sum of Z-scaled binding probabilities of paired epitopes that are unable to induce cross-reactivity (non-cross conserved epitopes) within two protein sequences using equations (1.1 and 1.2). *d* is the T cell immunity distance between a pair of epitopes, *i* and *j* are the non-cross conserved T cell epitopes from two protein sequences, *α* is a class I or class II allele, *p* is the predicted binding probability against allele *α*, A is a set of alleles.

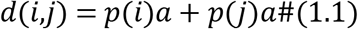

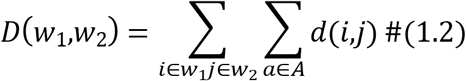

To evaluate the T cell epitope immune distance generated by the EpiCC algorithm, we further adapt the equations (1.1 and 1.2) to re-calculate T cell epitope immune distance with customized Python scripts using MHC binding prediction results that are generated from publicly available T cell epitope prediction tool, netMHCpan EL 4.1 methods in the Immune Epitope Database (IEDB) [51]. Eigenvalues of each sequence that were calculated from the pairwise distance matrix with “RSpectra” package were used to statistically examine the correlation of the epitope distances that are computed from the two methods, and Pearson correlation test was used to test the correlation hypothesis.

The cross-conservation of vaccine strains against circulating RSV can be evaluated by the cross-conservation of the epitopes within vaccine strains (*v*) and wild circulating strains (*w*). T cell cross-conservation between two epitopes can be represented by a joint probability estimation and therefore T cell cross-conservation between two sequences can be represented by summing T cell cross-conservation of the paired T-epitopes within two protein sequences. The proportion of T cell cross-conservation between the vaccine and circulating strains (*P*) with a set of alleles (A) can be represented as the equations (2.1 and 2.2), where *p* is the predicted binding probability in EpiMatrix, *i* and *j* are the cross conserved T cell epitopes, *α* is a class I or class II allele.

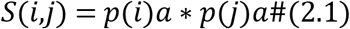

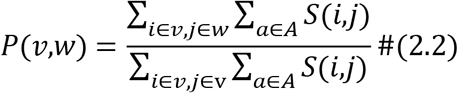

### Dimension reduction

The equation to calculate T cell epitope immune distance was applied iteratively to the subsampled dataset and therefore the pairwise T cell epitope immune distances are structured into an *n x n* square-distance matrix. Given that each protein is described by a relative distance to the rest of n-1 proteins, the data must be dimensionally reduced to be graphed. Classic (metric) multidimensional scaling (MDS) can be used to preserve the distances between a set of observations in a way that allows the distances to be represented in a two-dimensional space. MDS was performed as previously described by Gower [52]. The MDS method first constructs an n-dimensional Euclidean space using the distance matrix in which all distances are conserved, and then principal component analysis is performed. MDS and Goodness-of-fit (GOF) [53] were carried out using the *cmdscale* package in R [52]. K-means clustering was performed using the *kmeans* function in base R. Due to the lack of previous characterizations of RSV T cell immunity clusters, the number of T cell immunity groups was determined using the optimized within-cluster sum of square (wss) with Elbows method [54].

### Calculation of genetic hamming distance

Genetic hamming distance, which is defined as the number of bases by which two nucleotide sequences differ, was calculated by comparing the number of different bases between each sequence in the subsampled datasets. The reconstructed most recent common ancestor (TMRCA) sequences for each dataset (subsampled F and G protein sequences of subtype A and subtype B, respectively) were estimated using the program “Treetime” and were used as root in our analysis [45].

## Data and materials availability

Accession number to RSV sequence in the paper are available in supplementary materials. Code to generate T epitope landscapes are deposited in GitHub https://github.com/JianiC/RSV_Epitope.

## Supporting information

**S1 Fig. Distribution and diversity of T cell epitopes in RSV F protein.** The tree panel on the left is a time-scaled phylogeny build with RSV-A **(A)** or RSV-B **(B)** F gene nucleotide sequences using the ML approach. Determined genotypes are labeled on the right with black bars. Each color column on the right side represents the presence of an MHC class I or class II epitope. Only the epitopes that are present in more than 1% of sampled isolates are displayed. The column color indicates different numbers of epitope sequences at the same location.

**S2 Fig. Distribution and diversity of T cell epitopes in RSV G protein.** The tree panel on the left is a time-scaled phylogeny build with RSV-A **(A)** or RSV-B **(B)** G gene nucleotide sequences using the ML approach. The clades that contain novel 72-nt or 60-nt duplication at the second hypervariable region of G gene were highlighted in red. Determined genotypes are labeled on the right with black bars. Each color column on the right side represents the presence of an MHC class I or class II epitope. Only the epitopes that are present in more than 1% of sampled isolates were displayed. The column color indicates different numbers of epitope sequences at the same location.

**S3 Fig. Distribution of JanusMatrix Human Homology score for putative RSV MHC class I and class II epitopes.** The cross-reactive potential of identified putative T cell epitopes and human host was represented with a JanusMatrix Human Homology score. 6.45% identified putative class I epitopes and 1.12% class II epitopes are cross-conserved on the TCR face with human epitopes.

**S4 Fig. Predicted T cell epitope landscapes of RSV surface proteins.** RSV T cell epitope landscapes were built with sequenced-based MHC class I epitope binding prediction (left), MHC class II epitope binding prediction (middle) or combining class I and class II epitope biding prediction (right). Sequences are colored by the epitope cluster determined by epitope landscapes built with combining Class I and Class II epitope prediction

**S5 Fig. Total within sum of squares (*wss*) using *k-means* algorithm.** Totals within sum of squares in epitope topographies were calculated after clustering into k (from 1 to 10) groups with *k-means*. The optimal number of clusters is determined to be 3 in the analysis of RSV-A F and G proteins and is determined to be 2 in the analysis of RSV-B F and G proteins using the Elbow method.

**S6 Fig. Validation of T cell epitope distance estimation using the IEDB analysis resource**. Validation is performed with MHC class I epitope binding prediction of RSV-A F protein. **(A)** Heatmaps for pairwise MHC class I epitope distance estimated in iVAX toolkits or calculated with custom python scripts using MHC class I molecule binding prediction that is implemented in IEDB. **(B)** Eigenvalues for each sequence are calculated from pairwise distance matrices using “RSpectra” package in R. The Pearson correlation test significantly supports a non-zero correlation between T cell epitope distance estimated with EpiCC and T cell epitope distance estimated with IEDB. **(C)** T cell epitope topographies are built with pairwise epitope distances estimated from EpiCC or IEDB. Both methods resulted in a similar cluster pattern for the CD8 T cell epitope profile of RSV-A F protein.

**S7 Fig. Evaluation of RSV vaccine candidate strains with T cell epitope content in different WHO regions**. RSV-A and RSV-B major surface protein sequences were grouped by isolation year and 6 isolated WHO regions, African Region (AFRO), Region of the Americas (PAHO), South-East Asia Region (SEARO), European Region (EURO), Eastern Mediterranean Region (EMRO) and Western Pacific Region (WPRO). Each year group was labeled by the latest isolated year of sequences after the previous group label. The proportion of cross-conserved T cell epitope content between vaccine strains (CP248 or CP52) and wild circulating strains in different year groups was represented by bar graphs.

**S1 File. Accession number to RSV sequence that are used in this study.**

